# Structural basis for the increased processivity of D-family DNA polymerases in complex with PCNA

**DOI:** 10.1101/2020.01.29.925263

**Authors:** Clément Madru, Pierre Raia, Inès Hugonneau-Beaufet, Gérard Pehau-Arnaudet, Patrick England, Erik Lindahl, Marc Delarue, Marta Carroni, Ludovic Sauguet

## Abstract

Replicative DNA polymerases (DNAPs) have evolved the ability to copy the genome with high processivity and fidelity. In Eukarya and Archaea, the processivity of replicative DNAPs is greatly enhanced by its binding to the proliferative cell nuclear antigen (PCNA) that encircles the DNA. We determined the cryo-EM structure of the DNA-bound PolD-PCNA complex from *Pyrococcus abyssi* at 3.77Å. Using an integrative structural biology approach - combining cryo-EM, X-ray crystallography and protein-protein interaction measurements - we describe the molecular basis for the interaction and cooperativity between a replicative DNAP and PCNA with an unprecedented level of detail. PolD recruits PCNA *via* a complex mechanism, which requires two different PIP-boxes. We infer that the second PIP-box, which is shared with the eukaryotic Polα replicative DNAP, plays a dual role in binding either PCNA or primase, and could be a master switch between an initiation phase and a processive phase during replication.

## Introduction

DNA replication is one of the most important functions in living organisms and viruses. It ensures the integrity of the genome and the accurate transfer of genetic information. DNA polymerases (DNAPs) are the key enzymes of DNA replication and diverse DNA repair processes (Kornberg and Baker, 2005). Cellular organisms typically use multiple DNAPs, which have been grouped into different families based on their sequence alignments: PolA, PolB, PolC, PolD, PolX, PolY, and Reverse transcriptase (Braithwaite and Ito, 1993; Delarue et al., 1990). Genomic DNA replication is carried out by so-called replicative DNAPs, which have evolved to copy the genome with high processivity and fidelity (Raia et al., 2019a). The main replicative DNAPs from Eukarya are found in family B, from Bacteria in family C, and from Archaea in families B and D. Across every domain of life, polymerase holoenzyme accessory proteins play an integral role in achieving the extraordinary efficacy and accuracy of the replicative polymerase complex. These include a sliding clamp that encircles the DNA (Stukenberg et al., 1991) and greatly enhances the processivity (Indiani and O’Donnell, 2006). The bacterial sliding clamp is referred to as the β clamp, while the eukaryotic and archaeal sliding clamp protein is called the proliferative cell nuclear antigen (PCNA) (Bruck and O’Donnell, 2001). Clamps are constructed from either two (β) or three monomers (PCNA) to yield a ring composed of six domains, which share similar protein folds (Kong et al., 1992; Krishna et al., 1994).

In eukaryotes, PCNA helps to recruit replicative DNAPs δ (Pol δ) and ε (Pol ε) to the DNA and plays an essential function in cell proliferation; PCNA inhibition is therefore considered as a valuable anticancer strategy (Altieri and Kelman, 2018). In Archaea, PCNA has been shown to recruit replicative DNAPs of both B- and D-families, respectively named PolB and PolD (Henneke et al., 2005; Tori et al., 2007). Organisms within the archaeal domain of life possess a simplified version of the eukaryotic DNA replication machinery. The archaeal PCNA shares 25% identity with the human PCNA, and PolD, despite having the two-barrel fold of multi-subunit RNA polymerases for its catalytic domain, shares intriguing similarities with the three main multi-subunit eukaryotic replicative DNAPs: Polα, Polδ and Polε. In particular, the PolD DP1 subunit and the C-terminal domain of the DP2 subunit are homologous to the regulatory B-subunit and the C-terminal domain of the catalytic A-subunit, which are found in all eukaryotic replicative DNAPs (Raia et al., 2019b; Sauguet et al., 2016). PolD is an archaeal replicative DNA polymerase (Cann et al., 1998; Ishino et al., 1998), which is widely distributed among Archaea (except in crenarchaea) and has been shown to be essential for cell viability (Berquist et al., 2007; Birien et al., 2018; Cubonová et al., 2013)(Sarmiento et al., 2013). Similar to other replicative DNA polymerases, the activity of PolD is strongly stimulated through its interaction with PCNA (Castrec et al., 2009; Henneke et al., 2005; Tori et al., 2007). PCNA binding partners carry short motifs known as the PCNA-interacting protein-box (PIP-box), but sequence divergent motifs have been reported to bind to the same binding pocket (Boehm and Washington, 2016). While the PIP-boxes are the best known PCNA-interacting peptides, other motifs including RIR and MIP motifs have been reported (Dherin et al., 2009; Gueneau et al., 2013). Since the first structures of sliding clamps were determined, about one hundred structures have been reported, in their apo form, bound with DNA, or in complex with various PIP-boxes and other PCNA-interacting motifs (Prestel et al., 2019). However, the only structure of a full-length replicative DNAP bound with PCNA and DNA that has been reported to date is the *Pyrococcus furiosus* PolB-PCNA-DNA ternary complex, which was determined by negative-staining electron microscopy at 19 Å resolution (Mayanagi et al., 2011).

Here, we present the cryo-EM structure of the DNA-bound PolD-PCNA complex from *Pyrococcus abyssi* at 3.77 Å using an integrative structural biology approach, combining cryo-EM, X-ray crystallography and protein-protein interaction measurements. This structure unveils the molecular basis for the interaction and cooperativity between the whole replicative polymerase and PCNA with an unprecedented level of detail. PolD recruits PCNA *via* a complex mechanism, which requires two different PIP-box motifs, a C-terminal and an internal one that has never been characterized so far. We infer that the C-terminal PIP-box, which is shared with the eukaryotic Polα replicative DNAP, plays a dual role in binding either PCNA or primase, and could be a master switch between an initiation phase and a processive phase during replication.

## Results

### Overall architecture of the DNA-bound PolD-PCNA processive complex

The *P. abyssi* PolD processive complex was reconstituted by incubating PCNA with the PolD exonuclease-deficient variant (Palud et al., 2008) (DP1 H451A) in a 3:1 ratio, in the presence of an 18-mer primed DNA duplex with a 7-nucleotide overhang and a non-hydrolysable nucleotide analogue. The reconstituted complex (317 kDa) was vitrified and its structure was determined using single-particle cryo-EM. The map was solved at an average resolution of 3.77 Å (**Table S1**, **Figure S1, Figure S2**). The essential PolD and PCNA DNA-binding regions, as well as the DP1-DP2 and DP2-PCNA interface regions showed a higher resolution map at 3-3.5 Å (**Figure 1A**, **Figure S2**). In these regions, the density map of the DNA-bound PolD-PCNA complex was of sufficient quality to allow *de novo* building of the majority of the protein. The map includes several regions for which no atomic model was known before, such as regions neighbouring the active site and the DP1-DP2 interface. In the peripheral region of the complex, the DP2 KH domain, the DP1 OB domain and some regions of the PCNA were found to be flexible and the local resolution map ranged between 4.0 and 4.5 Å (**Figure S2**). In these regions, crystal structures of PolD DP1 (144-619) and DP2 (1-1050) individual subunits (Sauguet et al., 2016b), and the structure of the *P. abyssi* PCNA (from this study using X-ray crystallography at 2.3 Å resolution) were used in model building. DNA was docked into the cryo-EM map, guided by the density for the duplex region showing minor and major grooves as well as the unambiguous position of purines and pyrimidines (**Figure S3**). However, no obvious density for single-stranded DNA and the incoming nucleotide was observed in the DP2 active site.

**Figure 1:**
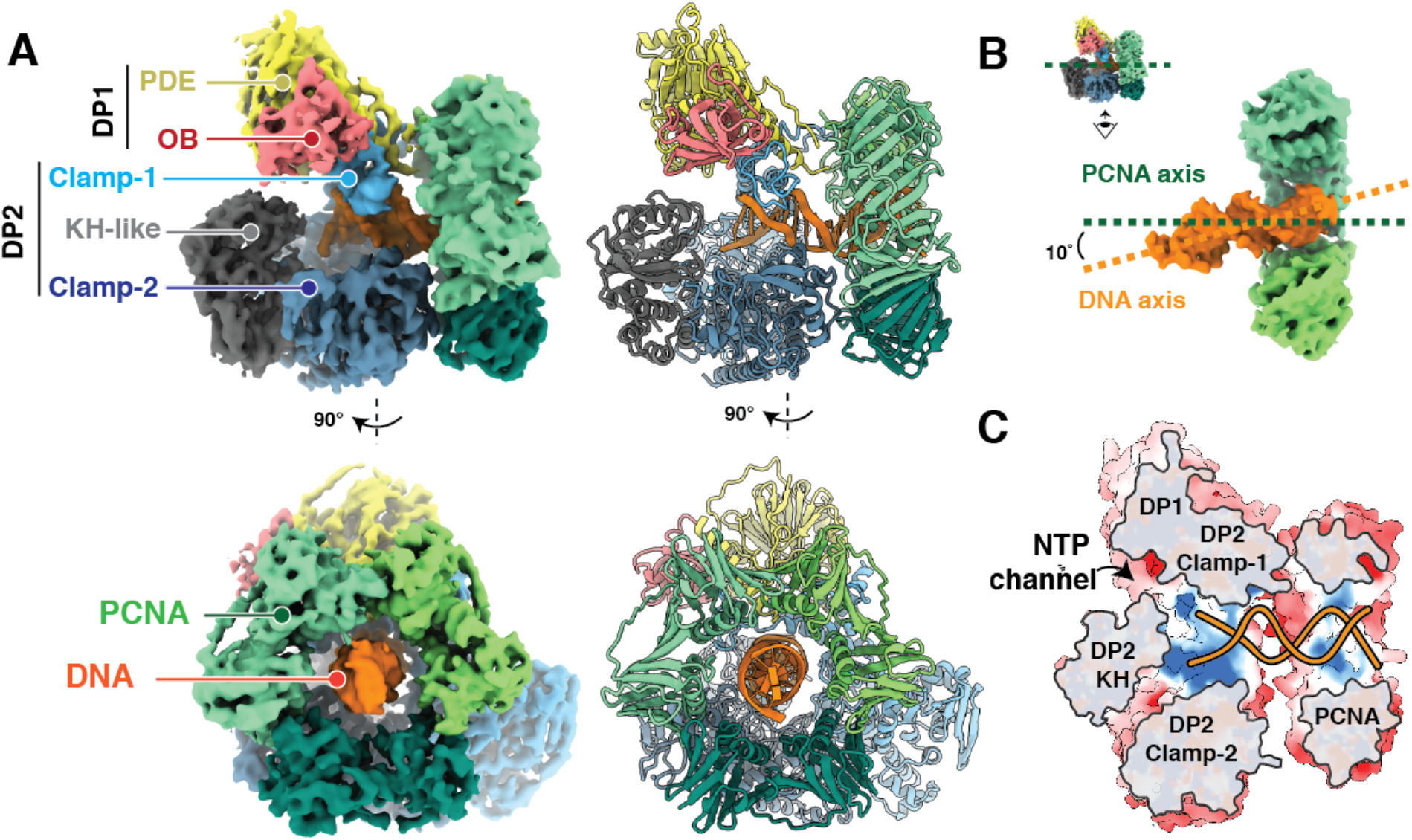
Cryo-EM structure of the DNA-bound PolD-PCNA processive complex. **(A)** Two orthogonal views of the cryo-EM density map (left) and cartoon representations (right) of the DNA-bound PolD-PCNA complex. **(B)** Orientation of the DNA duplex with respect to the PCNA threefold symmetry axis. **(C)** Cutaway front view of the PolD-PCNA-DNA complex showing the electrostatic surface potential with negative, neutral and positive charges represented in red, white and blue.

A defining feature of the PolD-PCNA-DNA ternary complex is its compactness: the radius of the PCNA ring and the clamp-like PolD DNA-binding domain match perfectly (**Figure 1A**). The structure of PCNA in the complex is not distorted compared to the structure of free PCNA and we conclude that the cryo-EM structure represents a stable interaction of DNA-bound PolD with a closed PCNA clamp. Away from the active site, PCNA surrounds one helix turn of the nascent DNA duplex, which is located at the centre of the PCNA ring. The nascent DNA duplex runs straight through PCNA and adopts an almost perpendicular orientation (~80°) with respect to the DNA (**Figure 1B**). In the 19 Å negative stain EM structure of *P. furiosus* PolB bound with PCNA (Mayanagi et al., 2011), and the 8 Å cryo-EM structure of the *E. coli* PolIIIα replicative complex bound with the sliding clamp β (Fernandez-Leiro et al., 2015), the DNA runs through the clamp in the same way. Interaction with PCNA nearly doubles the positively charged surface formed by the PolD active site, making a 60 Å long DNA binding site (**Figure 1C**). This structure thus rationalises how PCNA enhances the processivity of PolD: interaction with PCNA perpetuates the interactions with the nascent DNA duplex when it exits the PolD clamp, thereby preventing the polymerase from falling-off prematurely.

### PolD-DNA complex

The primer-template is held in position in the PolD active site by a bipartite clamp domain. Clamp-1 and clamp-2 domains contribute a central cleft located upstream from the DP2 catalytic centre, bordered with positively charged side chains, which encircles one helix turn of the nascent DNA (**Figure 2A**). While the DNA bound by the PolD-PCNA complex is predominantly in the B-form, interaction with the PolD clamp causes a distortion of the DNA region located next to the active site. Indeed, five base pairs at the primer 3’end are distorted, showing a decreased helical twist and a widened minor groove (**Figure 2B**). The clamp-1 domain contains a Zn-binding module, named Zn-III, connected to two α-helices that pushes against the minor groove of DNA. The Zn-III module harbours four conserved basic residues that interact intimately with the phosphodiester backbone: R1122 and K1129 interact with the primer strand while K1125 and K1145 contact the template strand. On the opposite side, the clamp-2 domain binds to the minor groove of the DNA, with numerous interactions between the side chains of five canonical lysines (K666, K668, K689, K785, K787) and both the primer and the template strands. A similar widening of the minor groove has been observed in the DNA-bound structures of other DNAPs associated with proofreading activities. In A-, B- and C-family DNAPs, conserved tyrosine, arginine or lysine residues have been shown to interact with the minor groove and to participate in the catalytic efficiency (Doublié and Zahn, 2014; Doublié et al., 1998; Wing et al., 2008). Minor groove hydrogen bonding interactions between DNAPs and N3 of purines or O2 of pyrimidines contribute to the efficiency of DNA synthesis and base selectivity. Similarly, the structure of PolD shows two canonical residues K1157 and Y1158 pointing towards universal hydrogen bond acceptors at purines N3 and pyrimidines O2 positions, which may be important for the catalytic efficiency and fidelity of PolD.

**Figure 2:**
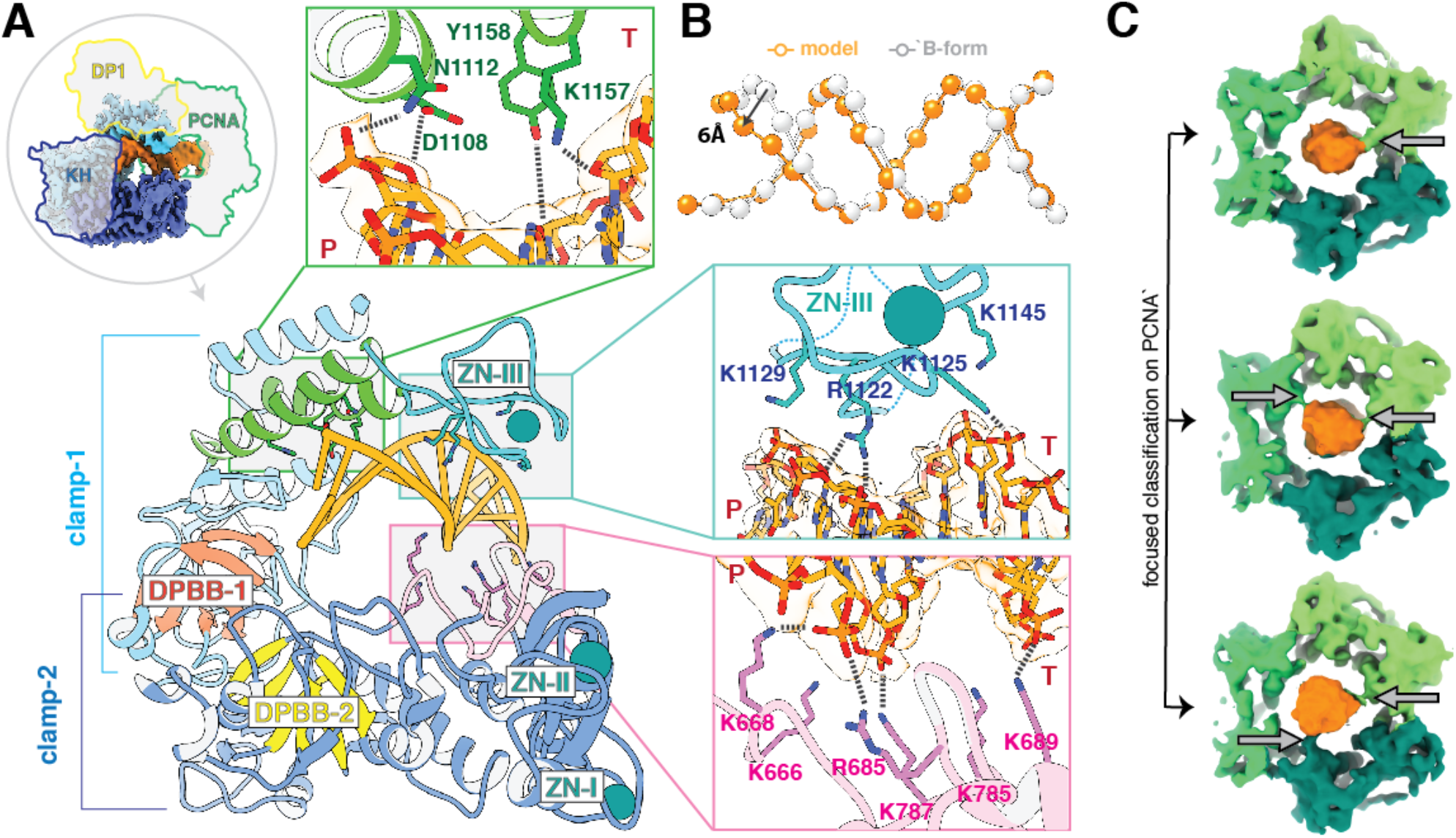
Structural basis for DNA binding by the PolD-PCNA complex. **(A)** View of the DP2 clamp-1 (light blue) and clamp-2 (dark blue) domains surrounding one turn of the DNA duplex (orange). The two catalytic double-psi β-barrels DPBB-1 and DPBB-2 are represented in red and yellow, respectively. Template and primer strands are indicated by T and P respectively. **(B)** Interaction with the PolD clamp causes a local distortion of the DNA (orange) compared with an ideal B-form DNA (white). The phosphates of the DNA duplexes are shown as spheres. **(C)** PCNA makes labile contacts with DNA. Focused classification on PCNA resulted in three 3D classes showing extra-densities between DNA and PCNA residues K84 and K86, which are underlined by arrows.

### PCNA-DNA binding interactions

Interestingly, PCNA and PolD adopt strikingly different DNA binding modes. While PolD extensively interacts with the nascent DNA duplex, the PCNA channel exposes several positively charged residues, which are pointed towards the DNA backbone and make labile transient polar contacts with DNA. Consistently, unlike the DNA region present in the PolD active site, the structure of DNA does not appear to be influenced by its interaction with PCNA and adopts a perfect B-form DNA architecture (**Figure 2B**). The local resolution in the cryo-EM map surrounding the PCNA is about 4-4.5 Å, substantially lower than the average resolution of the consensus map (3.77 Å), indicating a greater flexibility in the PCNA-DNA interactions compared to the PolD-DNA interactions. To characterise further the molecular determinants of the PCNA sliding movement, we performed a focused 3D-classification on PCNA and identified three classes showing extra-densities between DNA and the PCNA residues, K84 and K86 (**Figure 2C**). Interestingly, DNA was found to be in contact with different monomers of PCNA in the three 3D classes, suggesting that all three PCNA monomers contribute to DNA binding through short-lived polar contacts. Such labile transient interactions between DNA and PCNA have already been observed in another study using an integrative structural biology approach combining NMR and molecular dynamics simulations (March et al., 2017). PCNA thus provides an electrostatic cushion for the DNA to pass through as it leaves the PolD active site, thereby allowing it to rapidly slide onto DNA, pulled by PolD.

### PolD-PCNA interface

The cryo-EM map shows how PCNA is tethered to PolD through multiple contacts that involve both clamp-1 and clamp-2 domains of DP2 (**Figure 3A**). First, the α-helices, α-40 and α-41 of the clamp-1 domain are connected by a loop of 17 amino acids, which is hooked into PCNA. This loop binds to the canonical PCNA PIP-binding pocket through an internal PIP-box, which has never been identified so far (**Figure 3B**). Six residues within this internal PIP-box (hereafter referred to as iPIP) fill the PCNA PIP-binding pocket. Among them, the side chain of Q1198 penetrates deep into the pocket, making contacts with the canonical PCNA residue P245 (**Figure 3C**). In addition, the bulky sidechains of L1199, L1201 and I1202 make extensive contacts with hydrophobic residues, which line the PCNA PIP-binding pocket. Interestingly, the PolD iPIP shows a non-canonical structure compared to other PIP-boxes, lacking a four-residue 3_10_-helix-turn, which has been observed in the structures of all PIP-PCNA complexes determined so far (Prestel et al., 2019).

**Figure 3:**
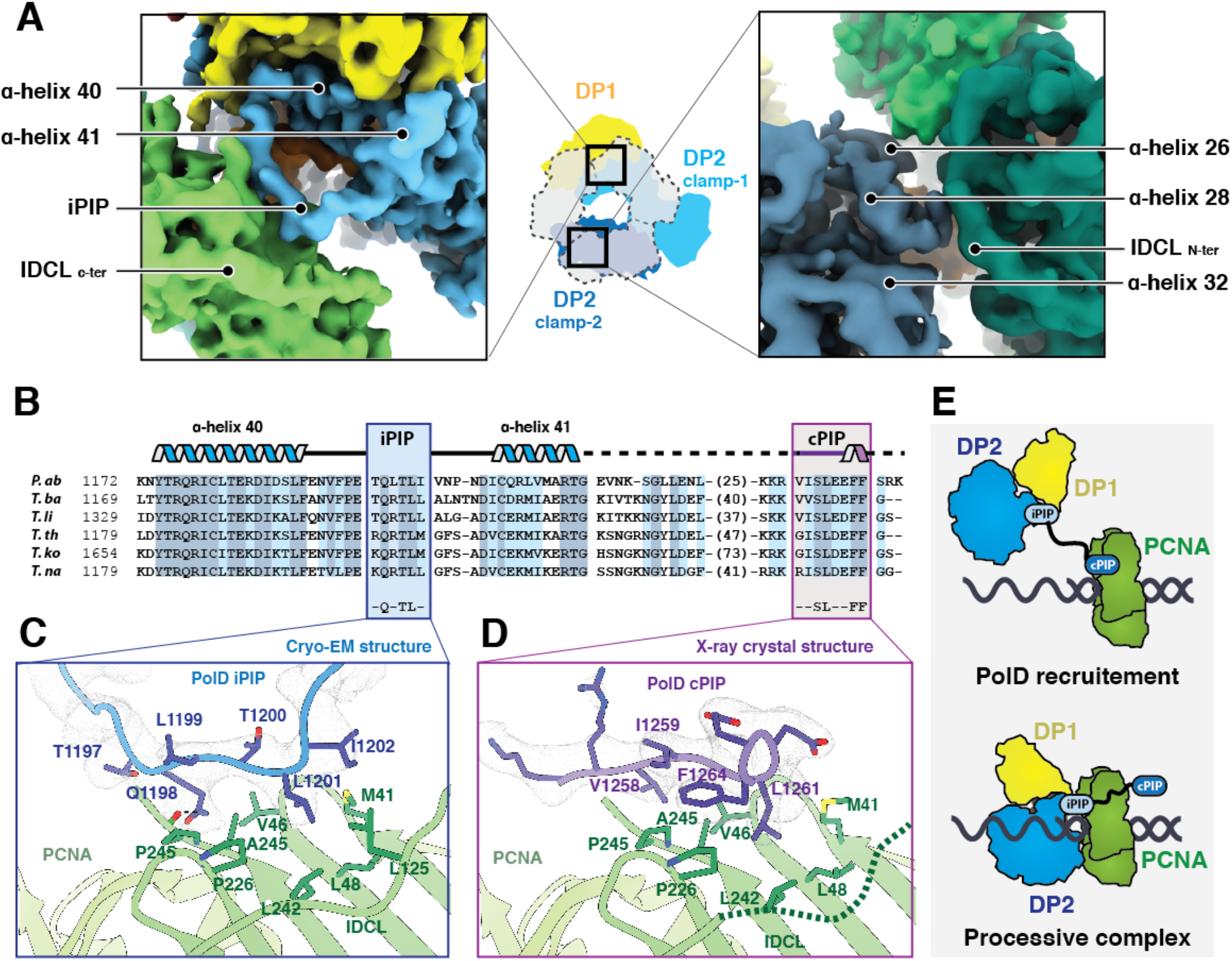
PolD uses two distinct PIP-boxes for molecular recognition of PCNA. **(A)** Two views of the cryo-EM map showing the interfacial region between the DP2 clamp-1 (left) and clamp-2 (right) binding to PCNA. DP2 clamp-1 and clamp-2 domains are represented in light and dark blue respectively, DP1 is shown in yellow, DNA in orange and PCNA in green. **(B)** Multiple-Sequence alignment of the C-terminal region in Thermococcus species: *Pyrococcus abyssi* (P.ab), *Thermococcus barophilus* (T.ba), *Thermococcus litoralis* (T.li), *Thermococcus thioreducens* (T.re), *Thermococcus kodakarencis* (T.ko), and *Thermococcus nautili (*T.na). Internal (iPIP) and C-terminal (cPIP) PIP-boxes are framed in blue and purple respectively, with secondary structure elements shown above. Sequence similarities are highlighted with light blue boxes and conserved residues are highlighted with dark blue boxes. **(C)** Detailed view of the iPIP-PCNA interaction in the cryo-EM map contoured at a level of 7 σ. **(D)** Detailed view of the cPIP-PCNA interaction in the 2Fo-Fc X-ray electron density map contoured at a level of 1.5 σ. **(E)** Hypothetical two-steps mechanism for PCNA recruitment by PolD.

Second, the N-terminal region of the interdomain-connecting loop (ICDL) of the adjacent PCNA monomer binds to the DP2 clamp-2 domain (**Figure 3A**). This interaction is mediated through polar contacts between residues E692, K779, Y781, and K896 from clamp-2 and PCNA residues H75, D117, and E119 (**Figure S4**). The large proportion of polar contacts underlines the plasticity of the latter interaction, which contrasts with the site-specific PCNA-clamp-1 interaction. This observation suggests that the PCNA-clamp-2 interaction may be easily broken or profoundly remodelled, enabling the PCNA ring to form a new interface with PolD, when the polymerase encounters a damage and adopts an editing mode. Such a conformational transition has been proposed for the PolB-PCNA complex, when PolB switches from the polymerase state to the proof-reading state (Mayanagi et al., 2011; Xu et al., 2016).

### PolD uses two-distinct PIP-boxes for molecular recognition of PCNA

In addition to the iPIP that was identified from our cryo-EM structure, the DP2 subunits of PolD from *P. furiosus* and *P. abyssi* have been shown to host a C-terminal PIP-box (Castrec et al., 2009; Tori et al., 2007). This second PIP-box (hereafter referred to as cPIP), is connected to DP2 by a 40-residue linker, which is variable in both length and amino acid composition across archaea (**Figure 3B**). Strikingly, both cPIP and the linker are not visible in the cryo-EM density of the DNA-bound PolD-PCNA complex. In order to better characterise the role of cPIP, we co-crystallized PCNA from *P. abyssi* with a 12 amino-acid peptide mimicking the DP2 cPIP and solved its structure at 2.7Å resolution (**Table S2 and Figure S5**). The final model includes 9 of the 12 amino-acids of the co-crystallized peptide. In contrast to the structure of iPIP, which differs from other structures of PIP-boxes, cPIP shares the same overall fold (**Figure 3D**) as those described in the literature (Prestel et al., 2019). Hence, the cPIP structure shows an extended peptide chain, whose C-terminal region folds into a 3_10_ helix. Several conserved hydrophobic residues of the cPIP - V1258, I1259, L1261, and F1264 - insert their bulky side chains into the hydrophobic cleft formed by the PIP-binding pocket on the PCNA surface. It is noteworthy that cPIP lacks the consensus Q residue, which is present in most PIP-boxes (Prestel et al., 2019).

The cryo-EM and crystal structures reveal that cPIP and iPIP adopt redundant binding positions in the PCNA PIP-binding pocket. Indeed, the side chains of L1199 and L1201 in iPIP and the side chains of I1259 and L1261 in cPIP are accommodated similarly in the PCNA-binding pocket, suggesting that the binding of iPIP and cPIP to PCNA are mutually exclusive. One may ask whether cPIP could not bind to one of the neighbouring PCNA subunits, but no extra density that could be accounted for by cPIP was found in the cryo-EM map. Furthermore, the linker-region connecting cPIP to DP2 can hardly cover the 70 Å distance, which separates the last defined residue of DP2 from the nearest unoccupied PIP-binding pocket. Altogether, these observations strongly suggest that iPIP and cPIP interact with PCNA through different mechanisms.

### The C-terminal PIP-box of PolD hosts a dual PCNA/primase binding-peptide shared with eukaryotic Polα

PolD shares, with its eukaryotic counterparts, unifying features of their subunit organisation that reveal a clear evolutionary relationship. The eukaryotic replicative DNAPs Polα, Polδ, and Polε possess a catalytic subunit, often referred to as the A-subunit, constitutively associated with different cohorts of regulatory proteins among which the B-subunits that are present in all three DNAPs assemblies (Doublié and Zahn, 2014). Both OB and PDE domains of DP1 share a remarkable degree of three-dimensional structural similarity with the regulatory B-subunits of all eukaryotic replicative DNAPs (Aravind and Koonin, 1998; Sauguet et al., 2016). In addition, the C-terminal region of their catalytic subunits, which is dedicated to interaction with their B-subunits, resembles the C-terminal region of DP2 that is required for interaction with DP1 (Raia et al., 2019b). In addition to their structural similarities (**Figure 5A**), PolD shares common functional features with Polα, which is tightly associated with the DNA primase in a complex called primosome that is required for initiating DNA replication in eukaryotic cells (Muzi-Falconi et al., 2003). Similarly, PolD has been shown to interact with the DNA primase (Pluchon et al., 2013) and is able to extend RNA primers (Henneke et al., 2005), suggesting that PolD is required for initiating DNA replication in Archaea. Previously, a short conserved motif located at the extreme C-terminus of Polα was shown to be critical for the interaction with the primase (Baranovskiy et al., 2016; Kilkenny et al., 2012, 2013). We tested whether the C-terminal region of PolD, which is homologous to that of Polα (**Figure 5B**), could host a similar primase-interacting peptide.

To assess the role of the C-terminus of PolD in the interactions with PCNA and primase, we performed biolayer interferometry (BLI) experiments using His_6_-tagged MBP-fusions of the C-terminal region of DP2, which were captured *via* surface-linked Ni-NTA. As expected, the MBP-iPIP-cPIP fusion (DP2:1196-1270) was found to readily bind to PCNA, with a K_D_ of 472±120 nM (**Figure 4A**). Interestingly, the same construct was also able to interact with primase, with a measured K_D_ of 237±22 nM (**Figure 4D**), which is very similar to the K_D_ of 245 nM that was reported for the interaction within the C-terminus of yeast Polα and the primase (Kilkenny et al., 2012). Deleting cPIP in the MBP-iPIP-ΔcPIP fusion (DP2:1196-1253), strongly impaired binding to PCNA and abrogated binding to the DNA primase (**Figure 4B, 4E**), showing that cPIP is a dual PCNA/primase binding peptide. Using His_6_-tagged PCNA and His_6_-tagged primase captured *via* surface-linked Ni-NTA, we then performed surface plasmon experiments, allowing us to measure the ability of PCNA and primase to bind a synthetic peptide encompassing the cPIP. The cPIP binds to PCNA with a lower affinity (K_D_ of 49±4 μM), which differs by two orders of magnitude from the one observed for the MBP-iPIP-cPIP fusion (**Figure 4C**), suggesting that the flanking region may be important for binding to PCNA. Consistently, the affinity of the PIP-box for PCNA can be modulated over four orders of magnitude by positive charges in the flanking regions (Prestel et al., 2019). Interestingly, cPIP binds to the primase substantially better than to PCNA, with a K_D_ of 4.0±1 μM (**Figure 4F**). The ability of cPIP to recruit both PCNA and primase is consistent with the dual role of PolD in DNA replication initiation and elongation, which requires interaction with both partners.

**Figure 4:**
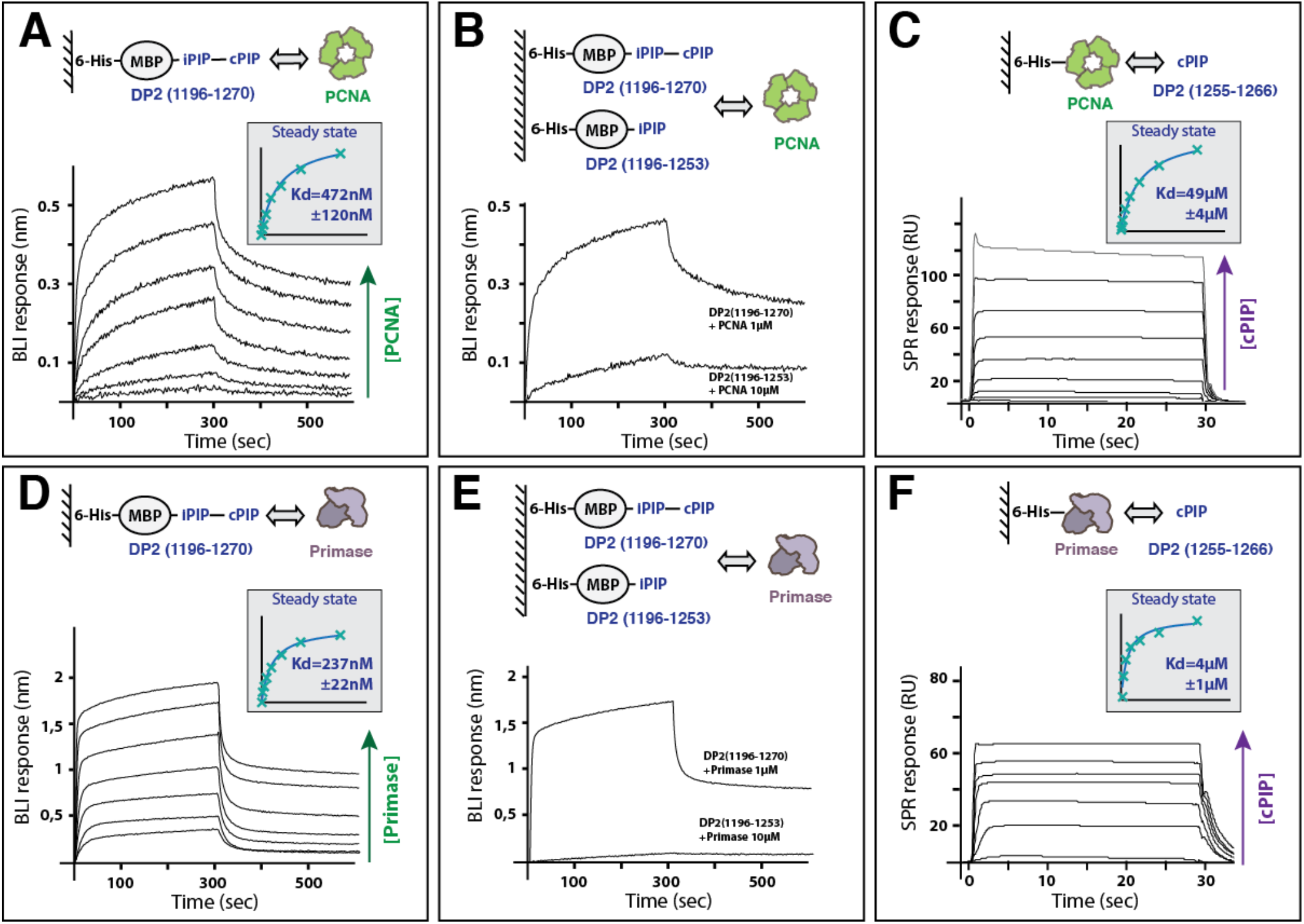
The C-terminal region of DP2 hosts a dual PCNA/Primase binding motif. **(A and D)** Specific binding of immobilized-DP2(1196-1270) to PCNA **(A)** and primase **(D)**, measured by BioLayer Interferometry (BLI). Steady-state analysis was performed using the average signal measured at the end of the association step (between 290 and 300s). **(B and E)** Comparative binding of immobilized-DP2(1196-1270) and -DP2(1196-1253) to PCNA **(B)** or primase **(E)** measured by BLI **(B)**. **(C and F)** Specific binding of immobilized-PCNA **(C)** and -primase **(F)** with increasing concentrations of cPIP by Surface Plasmon Resonance (RU: resonance units). Steady-state analysis was performed using the average signal measured at the end of the association step. The range of concentrations used in the binding experiments are listed in the method section.

**Figure 5:**
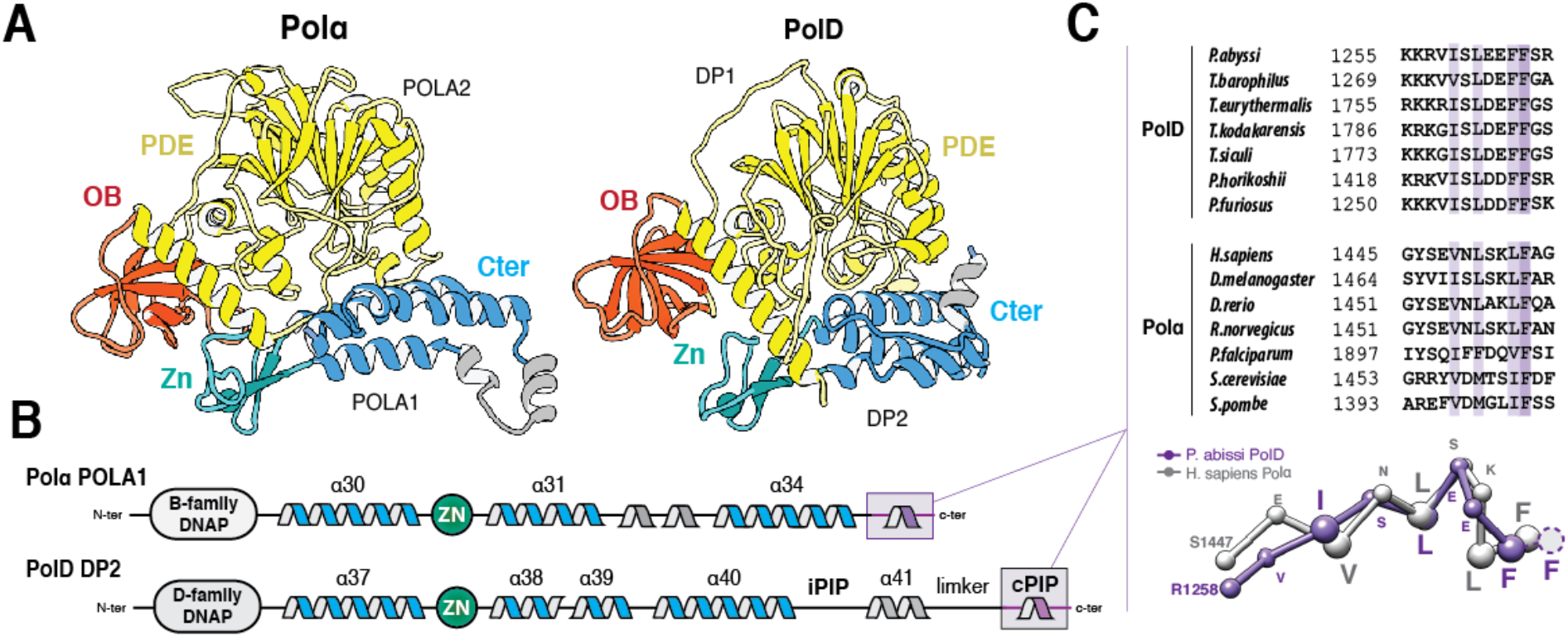
Shared primase binding peptide in archaeal PolD and eukaryotic Polα. **(A)** Structural comparison of the *P. abyssi* PolD DP1-DP2(1093-1216) region of the cryo-EM structure with the *H. sapiens* Polα POLA2-POLA1(1319-1456) crystal structure (PDB ID: 5EXR). **(B)** Shared structural features between archaeal PolD and eukaryotic Polα C-terminal regions. Conserved α-helices and Zn-binding are shown in blue and green, respectively. **(C)** The cPIP of DP2 resembles the primase-interacting motif located in the C-terminus of Polα. Top panel: Multiple-sequence alignment highlighting the conservation between the PolD cPIP motifs from Thermococcus species and the primase-interacting peptides of Polα. Sequence similarities are highlighted with light purple boxes, while conserved residues are shown with dark purple boxes. Bottom panel: Superimposition of the X-ray crystal structures of the cPIP from *P. abyssi* and the primase-interacting peptide of *H. sapiens* Polα (PDB ID: 5EXR). Cα-traces are represented as ball and sticks. Conserved residues are highlighted using larger spheres.

Interestingly, comparing the structures of the cPIP of PolD and the primase-binding motif of Polα reveals that both peptides fold in a one-turn 3_10_ helix (**Figure 5C**). This structural similarity is underpinned by the conservation of four hydrophobic and aromatic residues. The side chains of these four conserved hydrophobic residues become buried at the hydrophobic protein-peptide interface in both PolD-PCNA (**Figure 3D**) and Polα-primase (Baranovskiy et al., 2016; Kilkenny et al., 2013). Moreover, hot-spot residues that were shown to be important for primase binding (Kilkenny et al., 2012) such as L1451 and F1455, are particularly well-conserved in archaea (**Figure 5C**). In archaea and eukaryotes, the primase forms a heterodimer composed of a small PriS subunit with the polymerase activity and a larger regulatory PriL subunit. The Polα primase-binding peptide binds onto a hydrophobic edge of the PriL subunit (Baranovskiy et al., 2016; Kilkenny et al., 2013). Interestingly, in Archaeal PriL, the exposed hydrophobic surface is buried by a short α-helix, which fills the space occupied by the DNAP in the Polα-primase complex (Kilkenny et al., 2013). This suggests that either cPIP binds to another site in the Archaeal primase or that the corresponding region is remodelled upon cPIP binding. Such differences in the mode of interaction between the primase and these two polymerases may be accounted for by the fact that, while the eukaryotic Polα forms a stable and constitutive complex with primase, PolD and primase form only a transient complex.

## DISCUSSION

In contrast with cellular transcriptases and ribosomes, which evolved by accretion of complexity from a conserved catalytic core, it is striking that DNA replication was reinvented several times during evolution and that no replicative DNA polymerase family is universally conserved. Bacteria, Archaea, and Eukarya have evolved three distinct protein folds to replicate their genomes: i) the Polβ-like fold - found in bacterial Pol-III, ii) the Klenow-like fold - found both in archaeal/eukaryotic B-family DNAPs and in bacterial Pol-I, iii) the two-barrel fold - the third structural class of DNAPs that was recently unveiled with the structure of archaeal PolD (Sauguet, 2019; Sauguet et al., 2016). While structurally divergent, all replicative DNAPs share unifying features. Hence, across every domain of life, the extraordinary efficacy of the replicative polymerase complex is dependent on their interaction with sliding clamps, which encircle DNA and greatly enhance their processivity (Bruck and O’Donnell, 2001). The structure of the DNA-bound PolD-PCNA complex from *P. abyssi* unveils the molecular basis for the interaction and cooperativity between an entire replicative polymerase and PCNA with an unprecedented level of details.

Away from the PolD active site, PCNA surrounds one helix turn of the nascent DNA duplex, perpetuating the interactions with the nascent DNA duplex, thereby preventing the polymerase from falling-off prematurely. While PolD makes extensive and strong contacts with the DNA minor groove, PCNA contributes to DNA binding through short-lived polar contacts, which provides an electrostatic cushion for the DNA to pass through as it leaves the PolD active site, thereby allowing PCNA to rapidly slide onto DNA, pulled by PolD. Interestingly, the archaeal PCNA-PolD complex shares intriguing structural features with the bacterial PolIIIα replicative complex bound to the sliding clamp β (Fernandez-Leiro et al., 2015). The bacterial clamp β and the archaeo-eukaryotic PCNA are constructed from two or three monomers, respectively, which share similar protein folds (Kong et al., 1992; Krishna et al., 1994). In both structures, the nascent DNA duplex runs straight through PCNA and adopts an almost perpendicular orientation with respect to the DNA. While the bacterial sliding clamps are phylogenetically distantly related to their archaeal and eukaryotic counterparts, all replicative polymerases share an evolutionary conserved mechanism of interaction with their sliding-clamps. Despite belonging to two structurally distinct classes of DNA polymerases, archaeal PolD and bacterial PolIII share similar clamp binding peptides, which fit into hydrophobic pockets located at the surface of the clamp. Hence, the PolD(iPIP)-PCNA and PolIII-clamp β interfaces show a remarkable degree of three-dimensional similarity (**Figure S6**). The common architecture of their interface is underpinned by the conservation of several hydrophobic and polar residues, such as Q1198, L1199 and L1201 on the polymerase side, and L48, L242 and P245 on the sliding clamp side (**Figure S6**). This site-specific interaction between replicative polymerases and their sliding clamps is conserved in all domains of life, probably inherited from the last universal common ancestor (LUCA).

PIP-boxes often exist in multiple copies in DNA polymerases. Polη and Polδ have three PIP-boxes, which contribute differentially to distinct biological functions (Acharya et al., 2011; Masuda et al., 2015). We have shown that PolD uses two distinct PIP-boxes for molecular recognition of PCNA, which are located in the C-terminal region of their DP2 subunit. Strikingly, these two PIP-boxes contribute differentially to PCNA recruitment. We hypothesise that PolD may be recruited by PCNA through a two-step mechanism (**Figure 3E**). First, PCNA is recruited by PolD through its interaction with the DP2 cPIP. Once the PolD-PCNA complex is loaded on DNA, the complex is stabilised by an interaction between PCNA and iPIP, as observed in the cryo-EM structure, while cPIP becomes dispensable. This mechanism is supported by a former study on PolD from *P. abyssi*, showing that removing the cPIP did not disrupt the physical interaction with PCNA, when both partners are bound to DNA (Castrec et al., 2009).

In addition, we have shown that cPIP is not only important for recruiting PCNA but does also interact with the DNA primase, a key actor of the replisome. The interplay at the replisome in hyperthermophilic Archaea is of special interest as their DNA is exposed to elevated temperatures (up to 113°C), which promote increased level of DNA damage (Lindahl, 1993). It is striking that these archaeal species manage to maintain their genome, with a reduced repertoire of DNA polymerases. While human cells are known to contain at least 17 different DNAPs (Yang and Gao, 2018), the hyperthermophilic archaeon *P. abyssi* only possesses three distinct DNA polymerases: PolD, PolB and the DNA primase-polymerase. Recent gene deletion studies on hyperthermophylic Euryarchaea have demonstrated that only PolD is required for viability, suggesting that PolD is solely responsible for DNA replication, while PolB may be required for DNA repair (Birien et al., 2018; Cubonová et al., 2013; Kushida et al., 2019). It is noteworthy that the situation is different in Crenarchaea that do not possess PolD (Yan et al., 2017). Due to the multiple biological reactions required during DNA replication, PolD must be able to switch from one replication factor to another in a spatially and temporally regulated process. Indeed, our work shows that cPIP has overlapping specificities and is capable of binding both PCNA and primase. Hence, PolD must be able to interact with the primase during the initiation of DNA replication, and with PCNA to ensure processive extension of both leading and lagging strands. The versatility of cPIP may be instrumental in such process. This finding expands current views on PCNA interactions showing that PIP-boxes are a much broader class of motifs than initially thought, which form the network of interacting proteins responsible for DNA replication and repair (Boehm and Washington, 2016).

In eukaryotes, chromosomal replication is accomplished primarily by three distinct DNAPs, which play different roles in DNA replication: Polα, Polδ, and Polε. Polα is tightly associated with the primase in a constitutive complex, named the primosome, which is responsible for initiating DNA replication (Muzi-Falconi et al., 2003). Polδ and Polε have been shown, in a series of experiments, to be responsible for lagging and leading strand replication, respectively (Nick McElhinny et al., 2008; Pursell et al., 2007). While they have diverged to acquire specific biological activities, all these three polymerases share unifying structural features that they most probably inherited from a common ancestor with PolD (**Figure 6**) (Raia et al., 2019b). Both OB and PDE domains of DP1 share a remarkable degree of three-dimensional structural similarity with the regulatory B-subunits of all eukaryotic replicative DNAPs (Aravind and Koonin, 1998; Sauguet et al., 2016). In addition, the C-terminal region of their catalytic subunits, which is dedicated to interaction with their B-subunits resembles the C-terminal region of DP2 that is required for interaction with DP1 (Raia et al., 2019b; Tahirov et al., 2009). Using an integrative structural biology approach, we identify here a conserved primase interacting peptide conserved in PolD and Polα. This finding extends the structural similarities between the archaeal and eukaryotic multi-subunit replicative DNAPs, suggesting that their common ancestor was associated with the primase. However, unlike the C-terminal region of PolD, which contains two PIP-boxes, the C-termini of Polδ and Polε contain no PCNA-interacting motif. We hypothesize that eukaryotic DNAPs evolved distinct mechanisms for recruiting PCNA, when the two-barrel D-family catalytic core found in PolD was exchanged by a Klenow-like B-family catalytic core, which is found in all contemporary eukaryotic replicative DNAPs. Interestingly, the recent cryo-EM structure of the *S. cerevisiae* Polδ (Jain et al., 2019) shows that the nascent DNA duplex is located about 52 Å away from the C-terminus of the polymerase (**Figure S7**), rationalizing why PCNA must occupy a distinct position in Polδ, compared with PolD. Altogether, elucidating the structure of the PolD-PCNA DNA-bound complex clarifies the evolutionary relationships with its eukaryotic counterparts and sheds light on the domain acquisition and exchange mechanism that occurred during the evolution, from the simpler replisome that prevailed in the last common eukaryotic-archaeal ancestor, to the more complex eukaryotic one.

**Figure 6:**
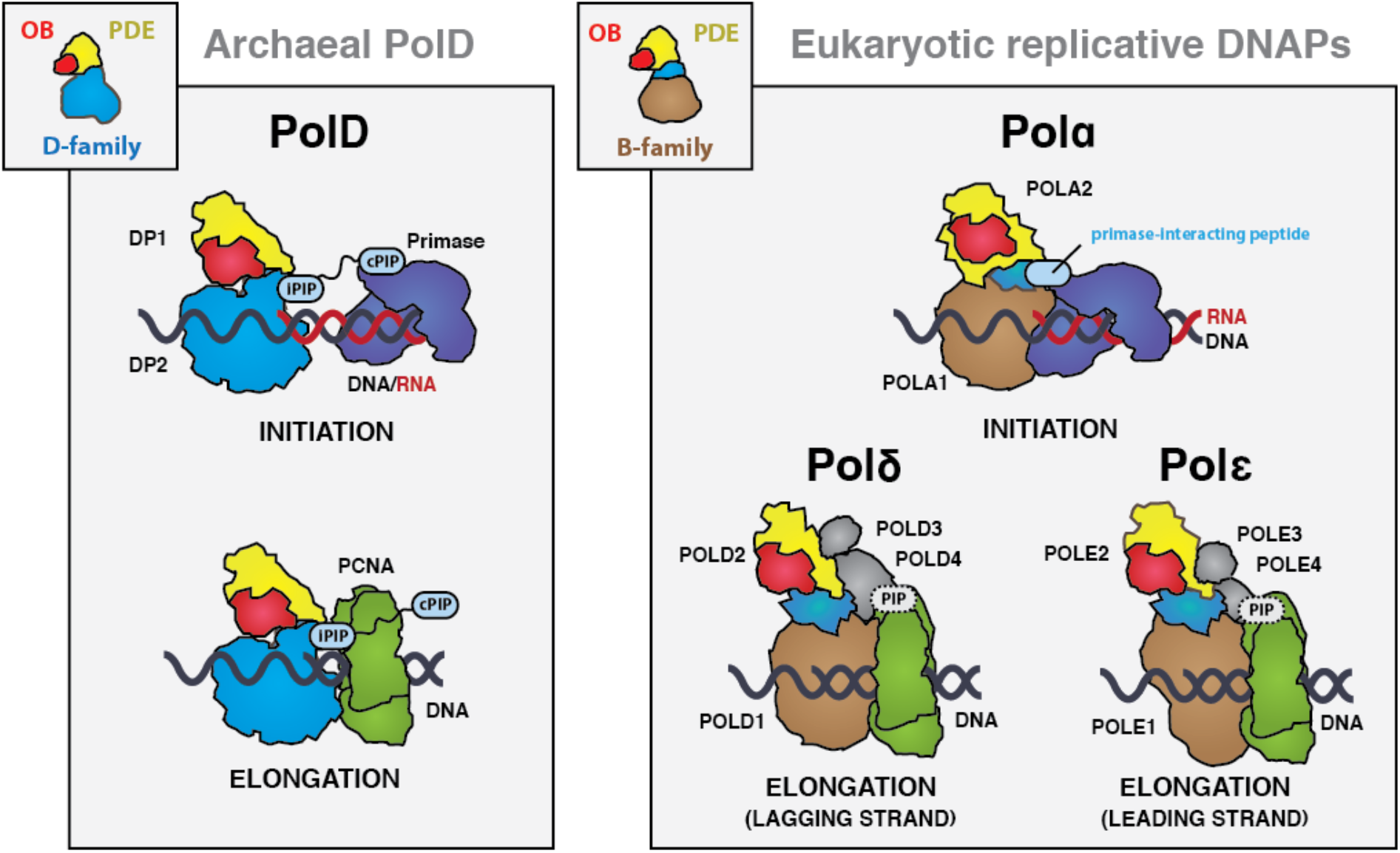
Comparison of multi-subunit polymerases in Archaea and Eukarya. The binding modes of PCNA and primase are illustrated in a schematic way as well as the two PIP motifs (cPIP and iPIP). The schematic representations are derived from the structures of the human Polα-primase complex (PDB ID: 5EXR) and the *P. abyssi* PolD-PCNA complex. No structures are currently available for the PCNA-bound Polδ and Polε complexes, but their schematic representations are supported by the literature (Acharya et al., 2011; Boehm et al., 2016; Burgers, 1991). DP2 is in light blue, the eukaryotic catalytic subunit is in brown, OB-fold is in red, PDE is in yellow, PCNA is in green and primase is in purple.

## METHODS

### Cloning, protein expression and purification

PolD from *P. abyssi* was purified as described previously (Raia et al., 2019b). The DNA coding sequence of DP2(1196-1253) and DP2(1196-1270) were inserted in a pIVEX His-MBP-TEV plasmid allowing the expression of a TEV-cleavable N-terminal 6xHis-MBP tag. The ORF of the PCNA gene from *P. abyssi* was optimized and synthesized by GeneArt (Thermo Fisher) and inserted into pet28-a(+) plasmid with a Thrombine-cleavable N-terminal 6xHis tag. The ORFs of the PriS and PriL(1-210) genes from *P. abyssi* were also optimized and synthesized commercially by GeneArt (Thermo Fisher) and inserted into pRSFduet(+) as a polycistronic construct with a TEV-cleavable N-terminal 14-His tagged PriS fusion protein. Production and purification of PolD constructions, PCNA and PriS-PriL(1-210) were performed as follow : proteins were expressed in BL21-CodonPlus (DE3)-RIPL strain from *E. coli* (Agilent) at 37°C in LB medium supplemented with 100 μg.mL^−1^ of antibiotic (kanamycin or ampicillin, depending on the plasmid) and 25 μg.mL^−1^ chloramphenicol. Recombinant protein expression was induced by adding 1 mM isopropyl-β-D-1-thiogalactopyranoside. Cells were then incubated overnight at 20°C, harvested by centrifugation, resuspended in buffer A (50 mM Na-HEPES at pH 8, 500 mM NaCl, 20 mM imidazole) supplemented with complete EDTA-free protease inhibitors (Roche) and lysed with a Cell-Disruptor. Lysates were then heated for 10 min at 60°C (except for the two MBP-fused DP2 constructs) and loaded onto 5-mL HisTrap columns (GE Healthcare) connected to an ÄKTA purifier (GE Healthcare). Elution was performed using a linear gradient of imidazole (buffer B, 50 mM Na-HEPES at pH 8, 500 mM NaCl, 0.5 M imidazole). The protein fractions were then combined, dialyzed in buffer C (20 mM Na-HEPES pH 8, 0.1 M NaCl), loaded onto 5-ml Heparin HiTrap HP columns (GE Healthcare) and eluted with a linear gradient, by mixing buffer C with buffer D (20 mM Na-HEPES pH 8, 1 M NaCl). Purifications were finally polished using exclusion-size chromatography in buffer E (20 mM Na-Hepes pH 8, 0,15 M NaCl) on a Superdex 75 10/300 or Superdex 200 10/300 (GE Healthcare) depending of the purified protein molecular weight.

### Sample preparation for Cryo-EM

The DNA duplex was prepared by mixing equivalent molar amounts of primer (CGCCGGGCCGAGCCGTGC) and template (AGGTCGTGCACGGCTCGGCCCGGCG). The mix was then heated 2 minutes at 90°C and slowly cooled to room temperature. PolD-PCNA-DNA complexes were associated by mixing 0.4 μM PolD and 1.2 μM PCNA with 0.7 μM DNA in the presence of 0.1 mM dAMPCPP in buffer F (20 mM Na-Hepes pH 8, 0.01 mM NaCl, 2 mM Mg-Acetate). The mixture was incubated for 20 min at 4°C, and pipetted onto glow-discharged holey carbon cryo-EM grids (C-flat 2/2, 4Cu, 50). Grids were frozen in liquid ethane by using a Vitrobot Mark IV (ThermoFischer) at 100% humidity, 22°C temperature, blotting force 20, and blotting time of 4s.

### Cryo-EM data acquisition and image processing

Movies were collected using EPU software on a Titan Krios electron microscope (Thermo Fisher Scientific) operated at 300 kV, on a GatanK2 Summit direct electron detector coupled with a Bioquantum energy filter with 20 eV slit. The defocus range was between −0.5 and −3.5 μm, the pixel size was 0.83 Å/pixel and the total dose was ~40 electrons/Å^2^, distributed into 40 frames. Image pre-processing until 2D classification was performed during data acquisition using the Scipion package (de la Rosa-Trevín et al., 2016). Images were imported and movie frames were aligned using MotionCor2 (Zheng et al., 2017) with dose compensation applied. The contrast transfer function was estimated with Gctf (Zhang, 2016) and particles were automatically picked using the Xmipp supervised picker after training was provided to tell particle from no-particles over ~1000 manually picked molecules (Abrishami et al., 2013). Particles were extracted 2x binned into a 150×150 pixel box to perform 2D classification for cleaning on-the-fly on consecutive batches of 20.000 particles each. Around 700.000 particles were automatically picked from 4602 micrographs and after 2D classification around 270.000 particles were kept, after excluding those belonging to poorly-resolved 2D classes.

Three initial models were generated by *ab initio* reconstruction using stochastic gradient descent in Relion 3.0 (Zivanov et al., 2018). At this point, good particles were again extracted without binning on 300×300 pixel box size and 3D refinement was run using one of the initial models. After 3D refinement, a 3D classification with 3 classes and local search was performed in order to separate possible conformations. The classification resulted in two classes displaying sharp details and a third class with less-defined features. No obvious conformational differences were noticed and the 3D classes better resolved were combined and 3D refined to a final consensus map at 3.77Å (gold standard 0.143 FSC criterion) resolution from around 150.000 particles. CTF-refinement and Bayesian polishing were also carried on, but they did not improve the overall resolution nor the quality of the map.

In order to separate different conformational states, focused 3D classification upon signal subtraction was performed using masks to focus on the PCNA part of the replicative complex. 3D classification without alignment led to 5 classes allowing the identification of 3 different types of weak contacts between PCNA and DNA. Classification was done in Relion using both a T parameter of 4 or 100 with the intent to catch some more details at higher T value. The results were comparable.

A summary of the full workflow is provided in **Figure S1** & **Figure S2**.

### Building and refinement of cryo-EM model

The density map of the DNA-bound PolD-PCNA complex was of sufficient quality to allow de novo building of the majority of the protein in COOT (Emsley et al., 2010). In the peripheral region of the complex, the DP2 KH domain, the DP1 OB domain and some regions of the PCNA were found to be more flexible and the local resolution map ranged between 4.0-4.5 Å. In these regions, model building was guided by the crystal structures of PolD DP1 (144-619) and DP2 (1-1050) individual subunits (Sauguet et al., 2016b), and the structure of the *P. abyssi* PCNA, which was solved in this study using X-ray crystallography at 2.3 Å resolution (PDB IDs: 5IJL, 5IHE and 6T7X). Concerning the DNA building, an ideal B-form DNA duplex was docked in the density as a starting point. The initial model was then subjected to global real-space refinement program from the PHENIX suite (Adams et al., 2011) using secondary structure restraints. The refined model was further manually inspected and adjusted in COOT. The final model was validated with statistics from Ramachandran plots and MolProbity scores (Davis et al., 2007). All figures were prepared using UCSF Chimera (Pettersen et al., 2004) and UCSF Chimera X (Goddard et al., 2018).

### Crystallization, X-ray data collection, and processing

PCNA crystallization trials were performed at 18°C using the hanging drop vapor diffusion technique in 2μL drops (1:1 reservoir to protein ratio) equilibrated against 500 μL of reservoir solution. For the PCNA-cPIP complex, PCNA was pre-incubated 30 minutes with a 2-fold molar excess of cPIP (KKRVISLEEFFS) (Smart Bioscience) in buffer E prior crystallization trials. PCNA crystals were obtained in 20% PEG 400, 0.2 M CaCl2 and 0.1 M MES pH 5.5 with a PCNA solution at 5 mg.mL^−1^ while PCNA-cPIP complex crystals were obtained in 30% PEG 400, 0.2 M MgCl2 and 0.1 M BIS-TRIS pH 7.1 with a PCNA-cPIP complex solution at 10 mg.mL^−1^. The crystals were cryoprotected by soaking in a 1:1 paraffin:paratone oil mix.

X-ray data were collected at the European Synchrotron Radiation Facility (ESRF) on beamlines ID23 and ID29 and at the SOLEIL synchrotron on beamlines PX1 and PX2. Datasets were indexed using XDS (Kabsch, 2010), scaled and merged with Aimless (from the CCP4 program suite (Collaborative Computational Project 1994) (Winn et al., 2011) and corrected for anisotropy with the STARANISO server (staraniso.globalphasing.org). PCNA X-ray structure was solved by molecular replacement using the structure of PCNA from *P. furious* (PDB ID: 5AUJ). Molecular replacements were carried out with the Phaser program from Phenix (Adams et al., 2011) and subsequent rebuilding and refinement were achieved with COOT (Emsley et al., 2010) and BUSTER (Blanc et al., 2004). Coordinates and structure factors of the PCNA and cPIP-bound PCNA structures from *P. abyssi* were deposited in the Protein Data Bank under accession codes 6T7X and 6T7Y, respectively.

### Bio-layer interferometry assays

Biolayer interferometry (BLI) experiments were performed on an Octet RED384 instrument (ForteBio). His-MBP-fused DP2 constructs were captured at a 1.5 nm density on Ni-NTA biosensors. Binding to PCNA and PriS-PriL(1-210) proteins was monitored for 300 s at 25°C in buffer E supplemented with 0.2 mg.mL^−1^ BSA. Seven proteins concentrations were assayed (31.25, 62.5, 125, 250, 500, 1000, 2000 nM) and a buffer-only reference was subtracted from all curves. Affinities were determined by fitting the concentration-dependence of the experimental steady-state signals, using the Octet RED data analysis v11 software (ForteBio).

### Surface plasmon resonance assays

Surface plasmon resonance experiments were performed using a Biacore T200 instrument (GE Healthcare). All measurements were performed at 25°C in buffer E supplemented with 100 μM EDTA. A series S sensor chip NTA (GE Healthcare) was used to immobilize approximately 2000 RU of His-tagged PCNA and PriS-PriL(1-210) on two of the four flowcells, and His-tagged MBP on a third as a reference. Ten concentrations (0, 0.75, 1.5, 3, 6, 12, 25, 50, 100 and 200 μM) of cPIP-peptide (KKRVISLEEFFS) (Smart Bioscience) were injected for 30 s at 30 μL.min^−1^ over the three flowcells. The raw sensorgrams were processed by subtracting both the signals measured on the reference flowcell and the signals measured for blank injections. Corrected data were analysed with the BIA evaluation software (GE Healthcare), by fitting the concentration-dependence of the experimental steady-state signals.

## References

Abrishami, V., Zaldívar-Peraza, A., de la Rosa-Trevín, J.M., Vargas, J., Otón, J., Marabini, R., Shkolnisky, Y., Carazo, J.M., and Sorzano, C.O.S. (2013). A pattern matching approach to the automatic selection of particles from low-contrast electron micrographs. Bioinforma. Oxf. Engl. 29, 2460–2468.

Acharya, N., Klassen, R., Johnson, R.E., Prakash, L., and Prakash, S. (2011). PCNA binding domains in all three subunits of yeast DNA polymerase δ modulate its function in DNA replication. Proc. Natl. Acad. Sci. U. S. A. 108, 17927–17932.

Adams, P.D., Afonine, P.V., Bunkóczi, G., Chen, V.B., Echols, N., Headd, J.J., Hung, L.-W., Jain, S., Kapral, G.J., Grosse Kunstleve, R.W., et al. (2011). The Phenix software for automated determination of macromolecular structures. Methods San Diego Calif 55, 94–106.

Altieri, A.S., and Kelman, Z. (2018). DNA Sliding Clamps as Therapeutic Targets. Front. Mol. Biosci. 5.

Aravind, L., and Koonin, E.V. (1998). Phosphoesterase domains associated with DNA polymerases of diverse origins. Nucleic Acids Res. 26, 3746–3752.

Baranovskiy, A.G., Babayeva, N.D., Zhang, Y., Gu, J., Suwa, Y., Pavlov, Y.I., and Tahirov, T.H. (2016). Mechanism of Concerted RNA-DNA Primer Synthesis by the Human Primosome. J. Biol. Chem. 291, 10006–10020.

Berquist, B.R., DasSarma, P., and DasSarma, S. (2007). Essential and non-essential DNA replication genes in the model halophilic Archaeon, Halobacterium sp. NRC-1. BMC Genet. 8, 31.

Birien, T., Thiel, A., Henneke, G., Flament, D., Moalic, Y., and Jebbar, M. (2018). Development of an Effective 6-Methylpurine Counterselection Marker for Genetic Manipulation in Thermococcus barophilus. Genes 9.

Blanc, E., Roversi, P., Vonrhein, C., Flensburg, C., Lea, S.M., and Bricogne, G. (2004). Refinement of severely incomplete structures with maximum likelihood in BUSTER-TNT. Acta Crystallogr. D Biol. Crystallogr. 60, 2210–2221.

Braithwaite, D.K., and Ito, J. (1993). Compilation, alignment, and phylogenetic relationships of DNA polymerases. Nucleic Acids Res. 21, 787–802.

Bruck, I., and O’Donnell, M. (2001). The ring-type polymerase sliding clamp family. Genome Biol. 2, reviews3001.1–reviews3001.3.

Cann, I.K., Komori, K., Toh, H., Kanai, S., and Ishino, Y. (1998). A heterodimeric DNA polymerase: evidence that members of Euryarchaeota possess a distinct DNA polymerase. Proc. Natl. Acad. Sci. U. S. A. 95, 14250–14255.

Castrec, B., Rouillon, C., Henneke, G., Flament, D., Querellou, J., and Raffin, J.-P. (2009). Binding to PCNA in Euryarchaeal DNA Replication requires two PIP motifs for DNA polymerase D and one PIP motif for DNA polymerase B. J. Mol. Biol. 394, 209–218.

Cubonová, L., Richardson, T., Burkhart, B.W., Kelman, Z., Connolly, B.A., Reeve, J.N., and Santangelo, T.J. (2013). Archaeal DNA polymerase D but not DNA polymerase B is required for genome replication in Thermococcus kodakarensis. J. Bacteriol. 195, 2322–2328.

Davis, I.W., Leaver-Fay, A., Chen, V.B., Block, J.N., Kapral, G.J., Wang, X., Murray, L.W., Arendall, W.B., Snoeyink, J., Richardson, J.S., et al. (2007). MolProbity: all-atom contacts and structure validation for proteins and nucleic acids. Nucleic Acids Res. 35, W375–W383.

Delarue, M., Poch, O., Tordo, N., Moras, D., and Argos, P. (1990). An attempt to unify the structure of polymerases. Protein Eng. 3, 461–467.

Doublié, S., and Zahn, K.E. (2014). Structural insights into eukaryotic DNA replication. Front. Microbiol. 5.

Doublié, S., Tabor, S., Long, A.M., Richardson, C.C., and Ellenberger, T. (1998). Crystal structure of a bacteriophage T7 DNA replication complex at 2.2 A resolution. Nature 391, 251–258.

Emsley, P., Lohkamp, B., Scott, W.G., and Cowtan, K. (2010). Features and development of Coot. Acta Crystallogr. D Biol. Crystallogr. 66, 486–501.

Fernandez-Leiro, R., Conrad, J., Scheres, S.H., and Lamers, M.H. (2015). cryo-EM structures of the E. coli replicative DNA polymerase reveal its dynamic interactions with the DNA sliding clamp, exonuclease and τ. eLife 4, e11134.

Goddard, T.D., Huang, C.C., Meng, E.C., Pettersen, E.F., Couch, G.S., Morris, J.H., and Ferrin, T.E. (2018). UCSF ChimeraX: Meeting modern challenges in visualization and analysis. Protein Sci. Publ. Protein Soc. 27, 14–25.

Henneke, G., Flament, D., Hübscher, U., Querellou, J., and Raffin, J.-P. (2005). The hyperthermophilic euryarchaeota Pyrococcus abyssi likely requires the two DNA polymerases D and B for DNA replication. J. Mol. Biol. 350, 53–64.

Indiani, C., and O’Donnell, M. (2006). The replication clamp-loading machine at work in the three domains of life. Nat. Rev. Mol. Cell Biol. 7, 751–761.

Ishino, Y., Komori, K., Cann, I.K., and Koga, Y. (1998). A novel DNA polymerase family found in Archaea. J. Bacteriol. 180, 2232–2236.

Jain, R., Rice, W.J., Malik, R., Johnson, R.E., Prakash, L., Prakash, S., Ubarretxena-Belandia, I., and Aggarwal, A.K. (2019). Cryo-EM structure and dynamics of eukaryotic DNA polymerase δ holoenzyme. Nat. Struct. Mol. Biol. 26, 955–962.

Kabsch, W. (2010). Integration, scaling, space-group assignment and post-refinement. Acta Crystallogr. D Biol. Crystallogr. 66, 133–144.

Kilkenny, M.L., De Piccoli, G., Perera, R.L., Labib, K., and Pellegrini, L. (2012). A conserved motif in the C-terminal tail of DNA polymerase α tethers primase to the eukaryotic replisome. J. Biol. Chem. 287, 23740–23747.

Kilkenny, M.L., Longo, M.A., Perera, R.L., and Pellegrini, L. (2013). Structures of human primase reveal design of nucleotide elongation site and mode of Pol α tethering. Proc. Natl. Acad. Sci. U. S. A. 110, 15961–15966.

Kong, X.P., Onrust, R., O’Donnell, M., and Kuriyan, J. (1992). Three-dimensional structure of the beta subunit of E. coli DNA polymerase III holoenzyme: a sliding DNA clamp. Cell 69, 425–437.

Kornberg, A., and Baker, T.A. (2005). DNA Replication (University Science Books).

Krishna, T.S., Kong, X.P., Gary, S., Burgers, P.M., and Kuriyan, J. (1994). Crystal structure of the eukaryotic DNA polymerase processivity factor PCNA. Cell 79, 1233–1243.

Kushida, T., Narumi, I., Ishino, S., Ishino, Y., Fujiwara, S., Imanaka, T., and Higashibata, H. (2019). Pol B, a Family B DNA Polymerase, in Thermococcus kodakarensis is Important for DNA Repair, but not DNA Replication. Microbes Environ.

Lindahl, T. (1993). Instability and decay of the primary structure of DNA. Nature 362, 709–715.

March, M.D., Merino, N., Barrera-Vilarmau, S., Crehuet, R., Onesti, S., Blanco, F.J., and Biasio, A.D. (2017). Structural basis of human PCNA sliding on DNA. Nat. Commun. 8, 1–7.

Masuda, Y., Kanao, R., Kaji, K., Ohmori, H., Hanaoka, F., and Masutani, C. (2015). Different types of interaction between PCNA and PIP boxes contribute to distinct cellular functions of Y-family DNA polymerases. Nucleic Acids Res. 43, 7898–7910.

Mayanagi, K., Kiyonari, S., Nishida, H., Saito, M., Kohda, D., Ishino, Y., Shirai, T., and Morikawa, K. (2011). Architecture of the DNA polymerase B-proliferating cell nuclear antigen (PCNA)-DNA ternary complex. Proc. Natl. Acad. Sci. U. S. A. 108, 1845–1849.

Pettersen, E.F., Goddard, T.D., Huang, C.C., Couch, G.S., Greenblatt, D.M., Meng, E.C., and Ferrin, T.E. (2004). UCSF Chimera--a visualization system for exploratory research and analysis. J. Comput. Chem. 25, 1605–1612.

Pluchon, P.-F., Fouqueau, T., Crezé, C., Laurent, S., Briffotaux, J., Hogrel, G., Palud, A., Henneke, G., Godfroy, A., Hausner, W., et al. (2013). An Extended Network of Genomic Maintenance in the Archaeon Pyrococcus abyssi Highlights Unexpected Associations between Eucaryotic Homologs. PLOS ONE 8, e79707.

Prestel, A., Wichmann, N., Martins, J.M., Marabini, R., Kassem, N., Broendum, S.S., Otterlei, M., Nielsen, O., Willemoës, M., Ploug, M., et al. (2019). The PCNA interaction motifs revisited: thinking outside the PIP-box. Cell. Mol. Life Sci.

Raia, P., Delarue, M., and Sauguet, L. (2019a). An updated structural classification of replicative DNA polymerases. Biochem. Soc. Trans.

Raia, P., Carroni, M., Henry, E., Pehau-Arnaudet, G., Brûlé, S., Béguin, P., Henneke, G., Lindahl, E., Delarue, M., and Sauguet, L. (2019b). Structure of the DP1-DP2 PolD complex bound with DNA and its implications for the evolutionary history of DNA and RNA polymerases. PLoS Biol. 17, e3000122.

de la Rosa-Trevín, J.M., Quintana, A., Del Cano, L., Zaldívar, A., Foche, I., Gutiérrez, J., Gómez-Blanco, J., Burguet-Castell, J., Cuenca-Alba, J., Abrishami, V., et al. (2016). Scipion: A software framework toward integration, reproducibility and validation in 3D electron microscopy. J. Struct. Biol. 195, 93–99.

Sauguet, L. (2019). The Extended “Two-Barrel” Polymerases Superfamily: Structure, Function and Evolution. J. Mol. Biol.

Sauguet, L., Raia, P., Henneke, G., and Delarue, M. (2016). Shared active site architecture between archaeal PolD and multi-subunit RNA polymerases revealed by X-ray crystallography. Nat. Commun. 7, 12227.

Stukenberg, P.T., Studwell-Vaughan, P.S., and O’Donnell, M. (1991). Mechanism of the sliding beta-clamp of DNA polymerase III holoenzyme. J. Biol. Chem. 266, 11328–11334.

Tahirov, T.H., Makarova, K.S., Rogozin, I.B., Pavlov, Y.I., and Koonin, E.V. (2009). Evolution of DNA polymerases: an inactivated polymerase-exonuclease module in Pol ∊ and a chimeric origin of eukaryotic polymerases from two classes of archaeal ancestors. Biol. Direct 4, 11.

Tori, K., Kimizu, M., Ishino, S., and Ishino, Y. (2007). DNA polymerases BI and D from the hyperthermophilic archaeon Pyrococcus furiosus both bind to proliferating cell nuclear antigen with their C-terminal PIP-box motifs. J. Bacteriol. 189, 5652–5657.

Wing, R.A., Bailey, S., and Steitz, T.A. (2008). Insights into the replisome from the structure of a ternary complex of the DNA polymerase III alpha-subunit. J. Mol. Biol. 382, 859–869.

Winn, M.D., Ballard, C.C., Cowtan, K.D., Dodson, E.J., Emsley, P., Evans, P.R., Keegan, R.M., Krissinel, E.B., Leslie, A.G.W., McCoy, A., et al. (2011). Overview of the CCP4 suite and current developments. Acta Crystallogr. D Biol. Crystallogr. 67, 235–242.

Xu, X., Yan, C., Kossmann, B.R., and Ivanov, I. (2016). Secondary Interaction Interfaces with PCNA Control Conformational Switching of DNA Polymerase PolB from Polymerization to Editing. J. Phys. Chem. B 120, 8379–8388.

Yang, W., and Gao, Y. (2018). Translesion and Repair DNA Polymerases: Diverse Structure and Mechanism. Annu. Rev. Biochem. 87, 239–261.

Zhang, K. (2016). Gctf: Real-time CTF determination and correction. J. Struct. Biol. 193, 1–12.

Zheng, S.Q., Palovcak, E., Armache, J.-P., Verba, K.A., Cheng, Y., and Agard, D.A. (2017). MotionCor2: anisotropic correction of beam-induced motion for improved cryo-electron microscopy. Nat. Methods 14, 331–332.

Zivanov, J., Nakane, T., Forsberg, B.O., Kimanius, D., Hagen, W.J., Lindahl, E., and Scheres, S.H. (2018). New tools for automated high-resolution cryo-EM structure determination in RELION-3. eLife 7.

